# DMAP – Graphical representation of physical and genetic map correlation

**DOI:** 10.1101/090415

**Authors:** David M.A. Martin, Glenn Bryan

## Abstract

Next-generation sequencing approaches coupled with appropriate assembly software can provide draft genome sequences of complex organisms as a series of unordered contigs in a timely and cost effective manner. Likewise, high throughput mapping technologies such as DArT and SNP platforms can provide a high density of sequence-anchored markers with which high resolution genetic maps can be constructed. Visualising and interpreting these data requires a new generation of tools as the volume of data leads to considerable redundancy and information overload in graphical representation. DMAP provides a highly configurable visual representation of physical and genetic map correlation, reducing data representation to an aesthetically acceptable degree. It also calculates an optimal orientation for the ordered sequence contigs, highlighting markers that are anomalous and contigs which may be in erroneous positions. Output is as PDF, allowing subsequent refinement prior to print publication and vector based representation for online supplementary figures. The perl scripts have few dependencies and code is freely available under a creative commons license (CC-BY) from the author’s GitHub repository at http://github.com/davidmam/DMAP.git.

## Motivation

The development in recent years of massively parallel shotgun sequencing technologies has revolutionised genome sequencing. Obtaining a genome sequence for a lower organism is relatively straightforward. Higher organisms however continue to pose a challenge, despite the high levels of genome coverage possible using the latest technologies. Whilst the majority of the sequence can be assembled into contigs, the presence of repetitive regions in the genome results in a large number of individual contigs. Orientation and arrangement of these requires them to be anchored to a high resolution genetic map which can provide an ordering for the contigs and potentially orientation information as well. Some progress can be made using paired end reads from next generation sequencing or through physical maps built using libraries of DNA fragments cloned into bacterial vectors such as Bacterial Artificial Chromosomes or FOSMIDs. These however can only group adjacent contigs rather than ordering contigs for which no relationship information is available. Contig anchoring requires a sequence-locatable marker, ie a specific sequence that can be unambiguously located in the genomic sequence, often referred to as a sequence-tagged-site [1]. Historically genetic maps have been prepared with a limited number of markers, typically a few tens per chromosome. However, recent technological developments allow the large scale (in the thousands to tens of thousands scale) identification of sequence-locatable markers. For example, diversity arrays (DArT) technology is a proprietary technology that identifies point mutation differences leading to restriction fragment polymorphism[2]. No prior knowledge of the sequence is required to develop the informative marker set which can then be directly sequenced and the sequences located in the genome using classical sequence similarity search methods. Perhaps the most commonly employed high-throughput marker technology is based on use of single nucleotide polymorphism (SNPs). SNP genotyping is commonly performed through use of technologies such as Illumina’s Oligo Pooled Array (OPA) Golden Gate methodology [3] and can also provide thousands to tens of thousands of potential sequence-locatable markers(e.g. [4-5]). SNP identification can be performed either by reference to the identified sequence or through *de novo* assembly of transcript or other sequences. Between them, these two methods can provide a pool of thousands to tens of thousands of informative markers. In the case of the potato genome project which inspired the development of this tool, these technologies give rise to around 200 segregating markers per chromosome.

Representing these sequence-associated markers in their genomic context graphically is a challenging task. These data can be incorporated into on-line comparison tools such as CMAP [6] but generating attractive and legible print output from such a tool is very difficult. It is practically impossible to present legibly even a few hundred markers on a chromosome in a standard print publication and selecting appropriate representative markers to view manually is a time consuming and tedious process. Also, individual markers may be misplaced either through errors in genotype data collection, map construction, sequence assembly or sequence anchoring. A new semi-automated approach was required for visualisation and analysis of these data so the DMAP tool was created.

### Methodology overview

DMAP reads sequence-anchored marker data, genetic map data and physical molecule composition (contig size, name and location) from text files in common formats along with formatting information. It then presents several analyses for the end user. The first is a linear mapping between the genetic map and chromosome physical map. The second is a correlation plot between the physical and genetic coordinates with a best-fit curve representing the relationship between recombination rate and physical distance. Finally a table of results is presented which identifies potentially problematic markers or contigs. The output is a PDF document from which relevant portions can be readily extracted and, should it be desired, edited in appropriate software (such as Inkscape [7] or Adobe Ilustrator [8]) prior to publication.

To prepare the linear map, DMAP bins the data on both genetic and physical axes. The genetic axis typically is binned at the average resolution of the map (approximately 1/size of mapping population) and the physical axis at a level which will allow marker names to be represented legibly in the published figure. Whilst sensible defaults are chosen by the software, these parameters are user customisable. DMAP then draws a link for each marker from the genetic map representation to the exact physical location. Genetic bins containing markers are labelled withtheir map position. No labels are shown for empty bins. Marker name labels are laid out in two columns according to an algorithm that checks for adjacent bin occupancy and attempts to fit as many labels in as feasible. The degree of ‘crowding’ of labels is a customisable parameter. Label types and format information can be specified and at least one marker label of each type per bin is selected for display. A portion of such a linear map is shown in Figure 1. It is the case that such maps do not always give an accurate impression of the data as a small number of erroneous markers will not show co-linearity between the physical and genetic map with a disproportionate visual impact in comparison to the majority of markers which do show co-linearity.

**Figure 1.**
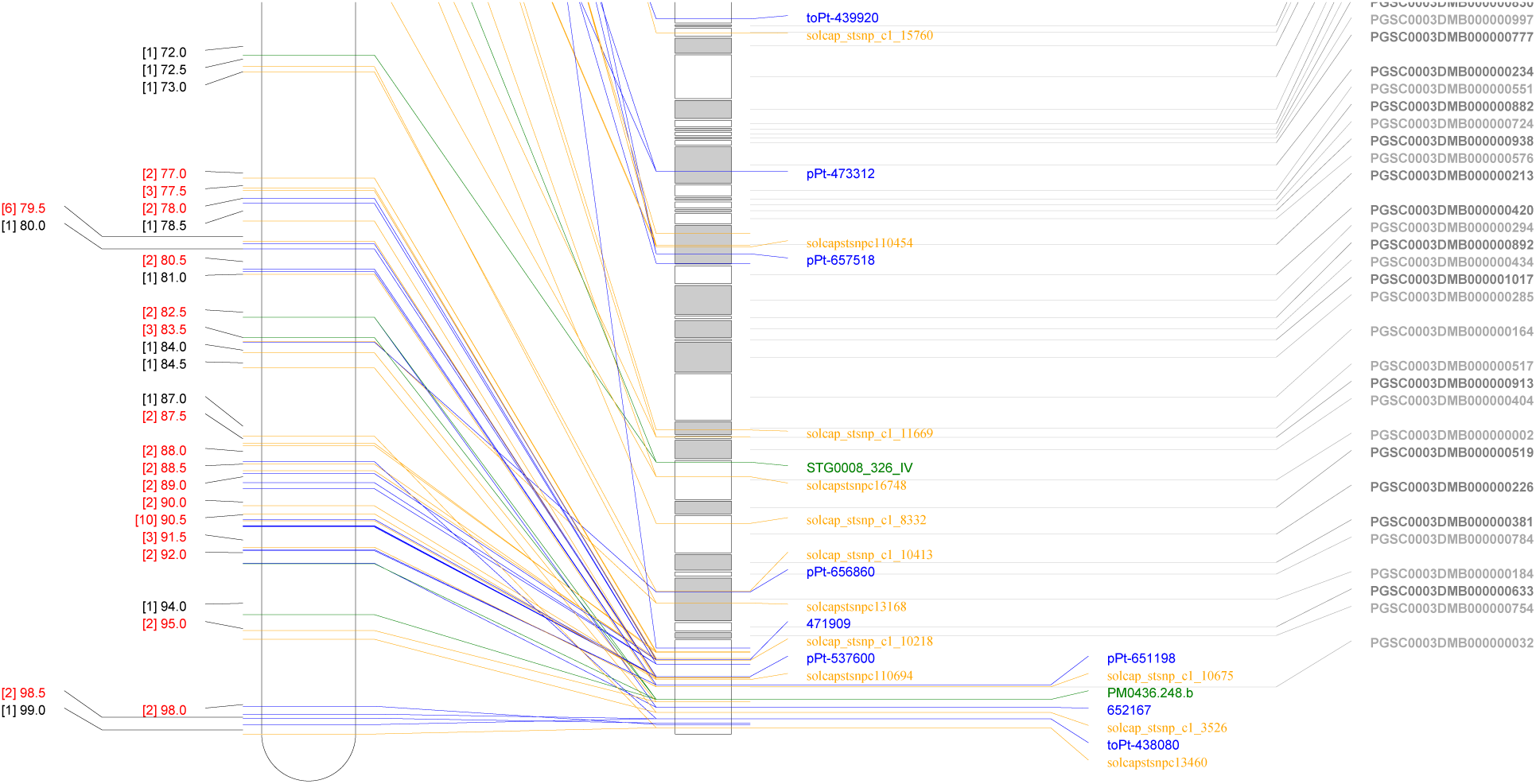
Colinear physical and genetic map of potato chromosome 4. A portion of a collinear map is shown. Links and text for markers are coloured according to the format described in the input file. Genetic marker bins are indicated with those containing more than one marker highlighted in red. Not all marker labels are shown and some are decanted to a second column automatically to improve readability. All molecular fragment labels are shown and coloured alternately grey and white for legibility.

To better explore the relationship between the physical and genetic maps, they are plotted on alternate axes (physical coordinate as the abscissa, genetic coordinate as the ordinate) (figure 2) and a hyperbolic sin curve is fitted to this data [equation 1] following the mapping function of Haldane [9]. The orientation of each molecule is optimised by testing the Chi-squared statistic for marker fit to the curve. Before and after curves and marker positioning are plotted on the figure, allowing immediate visual identification of potentially erroneous regions of the assembly, or artefcats in the genetic mapping data. The mapping function provides a good empirical fit but local conditions such suppression of recombination and recombination hotspots (e.g. [10,11]) will show deviations from the general trend.

**Figure 2.**
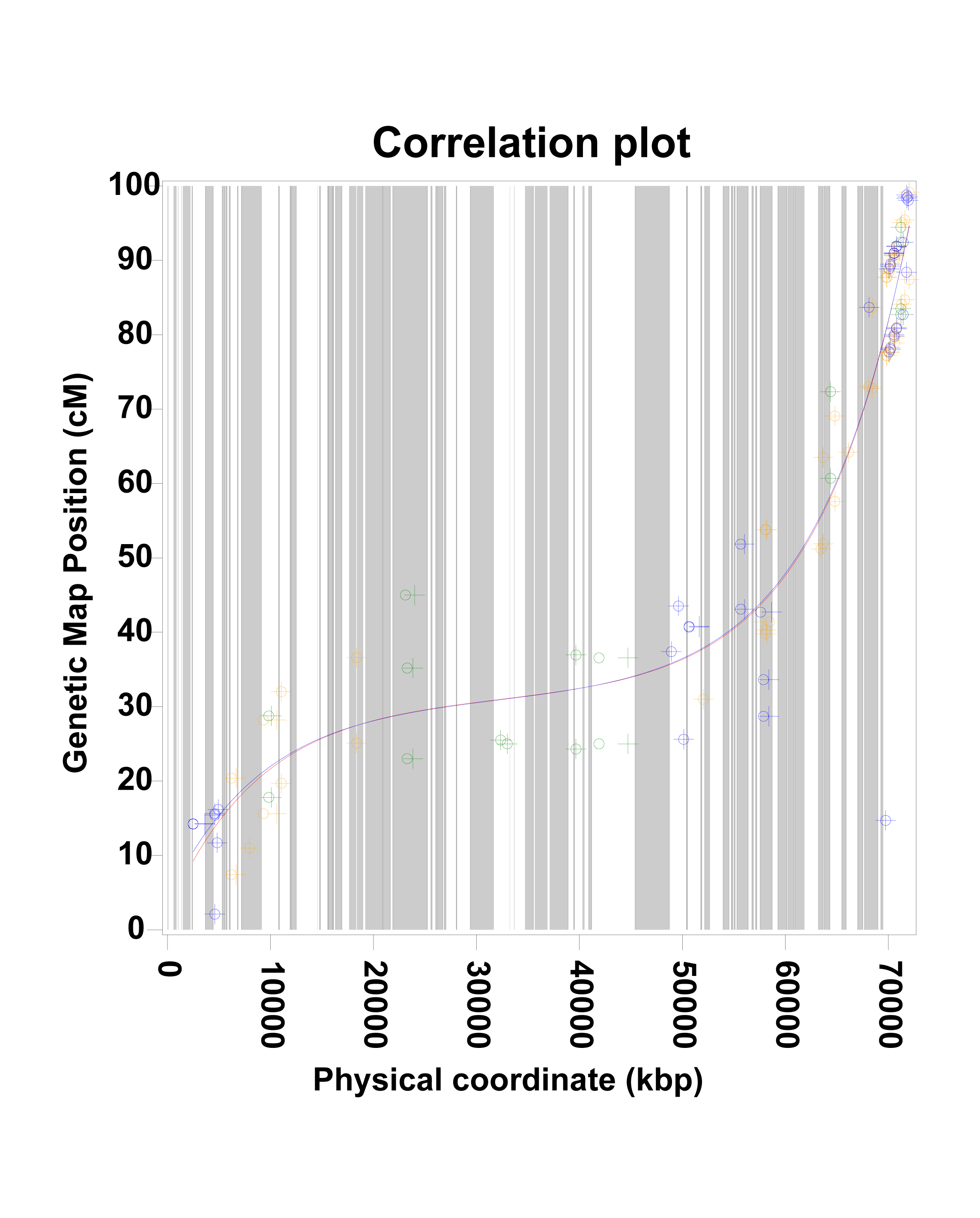
Correlation plot of genetic and physical coordinates. The observed data (+) is plotted against a background coloured to illustrate the individual molecule components of the physical coordinate (dark grey/light grey stripes). An initial curve fit to the observed data is shown (red) and a fit after reorienting of the molecule fragments has been performed (blue). Repositioned markers after reorientation are shown as circles.

### DMAP software implementation

DMAP is written in Perl as a command line application. It has few dependencies; the Perl Data Library (PDL) is required for curve fitting and the PDF::API2 library is required for PDF output. It should therefore run across most platforms though it has only been tested on Linux (x64) and Mac OS X. Runtime options are documented with perldoc within the code and in documentation included with the distribution. The software is available under a Creative Commons Attribution license and downloadable from the authors GitHub repository[12]

## Input format

DMAP reads input data from standard formats plus a display definition file. Pseudomolecule construction is read from an AGP file [13] specified by the –agpfile parameter which defines the order and size of each molecular fragment. Physical location for each marker is read from a GFF file [14] specified with the –gfffile parameter which links marker location, ID and type to a sequence coordinate. The ID and type are specified as key-value pairs in the note extended field (see example below.) Genetic map position is read from a JoinMap [15] output file specified with the –mapfile coordinate. An optional map name can be specified with the –mapname parameter. Thedisplay definition file is specified with the –infile option. It is a plain text file with one line per marker type and describes colour, font and line type for display of marker information. The file format for the display definition is shown in Table 1 and an example file fragment is shown in figure 3A.. The AGP file should contain a list of molecules and gaps. DMAP allows an AGP to represent a molecule as multiple dispersed fragments. Molecule names should be identical to those used in the GFF file and DMAP ensures that marker positions are mapped to the correct fragment. An example AGP file fragment is shown in figure 3B.

**Table 1:**
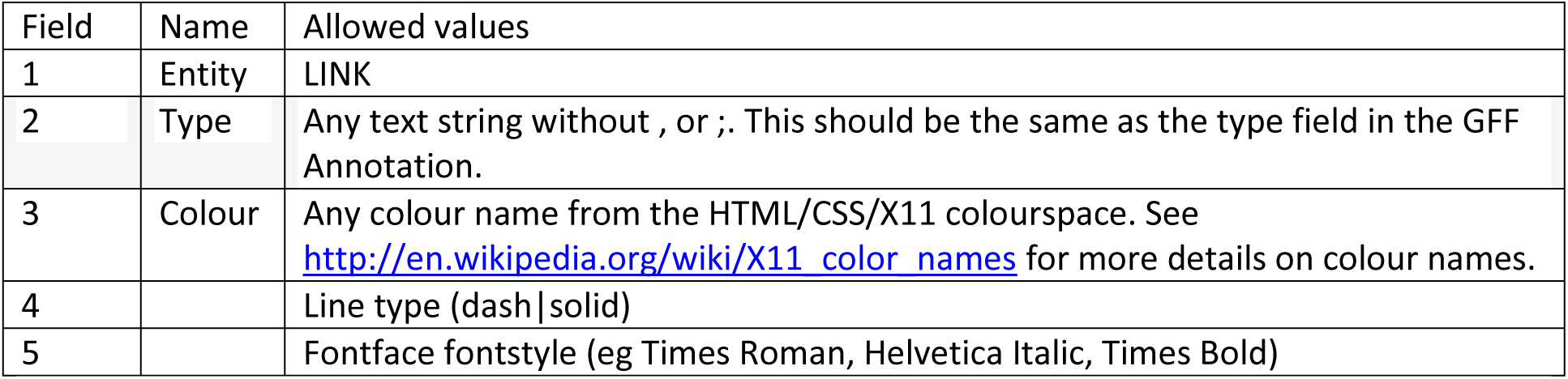
Syntax for display definition file. Each line is made up of comma separated fields. There should be no spaces next to the commas.

**Figure 3.**
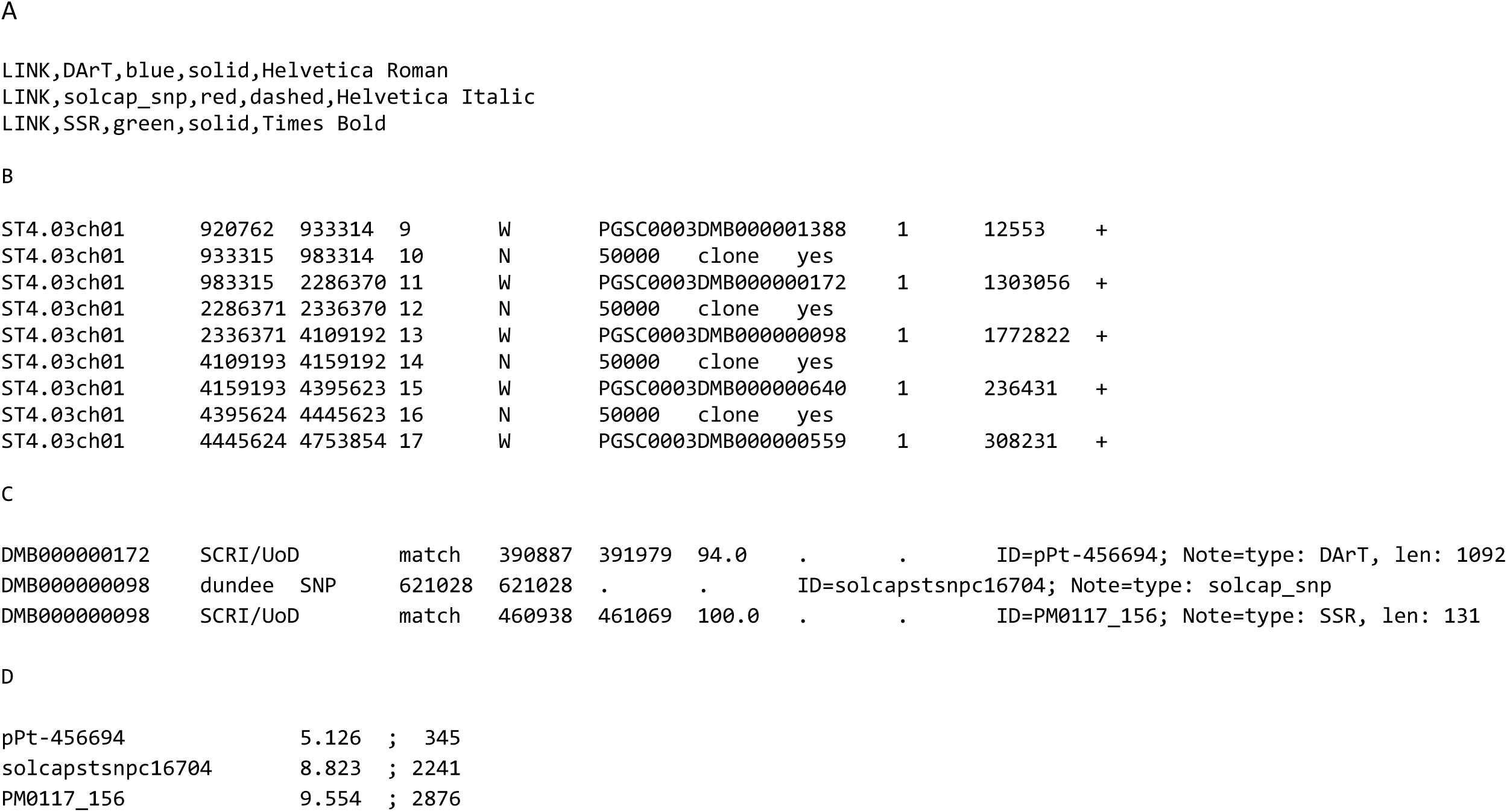
Example formats for data input to DMAP. A) Display Format specification file. B) AGP file describing pseudomolecule structure; C) GFF file linking marker positions to molecule fragments; D) Join Map output giving genetic distance for each marker

The required entries in the GFF annotation field are ID (which must be unique and identical to that used in the Joinmap file) and a value for ‘type’ in the Note annotation. The type value is used to determine formatting for the link and label. In the absence of a type specification, a default colouring scheme is used. If a molecule to which a marker has been mapped is not found in the AGP file, a warning message is output and that marker is ignored. The molecule name given in the first field must therefore match exactly the name used in the AGP file. An example GFF file fragment is shown in Figure 3C.

DMAP will read the standard Join Map output files. It requires that the marker name in the Join Map file is identical to that used in the ID annotation in the corresponding GFF file. If the marker ID is not located then that marker is skipped and a warning message issued. An example Join Map file fragment is shown in figure 3D.

### Results

DMAP produces a multi-page report in PDF format of which an example is shown in S1 in the supplementary material. The first page comprises a linear comparison of the genetic and physical map. A portion of the example file is shown in Figure 1. This is customisable through various command line parameters detailed in the documentation. The second page is a plot of genetic distance plotted against the physical distance. Each fragment is shown alternately as dark or light grey bands with markers plotted as a cross in the colour defined by their marker type. This plot enables rapid identification of markers that appear ‘out of place’ in the analysis, a factor that can be overemphasised in the linear map.

The third page is similar to page 2 but overlays the fitted relationship between genetic distance and physical distance (figure 2). In addition, the results of the orientation optimisation are illustrated with the markers displayed as circles. Fragments where the circles and crosses are not coincident will have been flipped in the optimised layout.

Finally DMAP reports the fit parameters (p1 to p4 from equation 1), the optimised fit parameters, the fit for each scaffold with the number of markers and the Chi2 value for the fit, a comparison between the physical map order for the marker-containing fragments and the genetic map order, and then a list of poorly fitting markers with the most discordant listed first.

### Conclusions

DMAP provides a simple and accessible method for visualisation and analysis of dense sequence-anchored genetic map data. It is simple to use and provides editor-friendly output suitable for direct use or downstream manipulation. In addition usage may be extended to compare genetic maps constructed from different varieties or strains where the markers are anchored to the same physical molecules, potentially adding valuable insight into the genomics of the organism.

## Supplementary-material

S1: Example DMAP report

## Equations

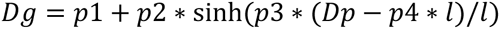

where *Dg* is the genetic map distance, *Dp* is the physical position of the marker, *l* is the chromosome length and *p1-p4* are optimsiable parameters with initial estimate values 50, 1, 5 and 0.5 respectively.

## References

[1] Olson, M., Hood, L., Cantor, C., Botstein D. A Common Language for Physical Mapping of the Human Genome. Science, 245:1434–5. 1989.

[2] JaccoudD, Peng K, Feinstein D, Kilian A (2001) Diversity Arrays: a solid state technology for sequence information independent genotyping. Nucleic Acids Research 29: e25.

[3] Fan JB, Chee MS, Gunderson KL. Highly parallel genomic assays. Nat Rev Genet. 2006

[4] Xuehui Huang, Qi Feng, Qian Qian, Qiang Zhao, Lu Wang, Ahong Wang, Jianping Guan, Danlin Fan, Qijun Weng, Tao Huang, Guojun Dong, Tao Sang, Bin Han High-throughput genotyping by whole-genome resequencing. Genome Research, Vol. 19, No. 6. (1 June 2009), pp. 1068–1076. doi:10.1101/gr.089516.108

[5] John Hamilton, Candice Hansey, Brett Whitty, Kevin Stoffel, Alicia Massa, Allen Van Deynze, Walter De Jong, David Douches, C. Robin Buell Single nucleotide polymorphism discovery in elite north american potato germplasm BMC Genomics, Vol. 12, No. 1. (9 June 2011), 302. doi:10.1186/1471–2164-12–302_Aug;7(8):632-44.

[6] Ken Youens-Clark, Ben Faga1, Immanuel V. Yap, Lincoln Stein and Doreen Ware (2009) CMap 1.01: a comparative mapping application for the Internet Bioinformatics (2009) 25 (22): 3040–3042.

[7] Inkscape drawing package [http://www.inkscape.org]

[8] Adobe Illustrator drawing program. [http://www.adobe.com]

[9] Haldane JBS. The combination of linkage values and the calculation of distances between the loci of linked factors. J Genet. 1919;8:299–309.

[10] Stirling, B., Newcombe, G., Vrebalov, J., Bosdet, I., Bradshaw Jr., H.D. 2001 Suppressed recombination around the MXC3 locus, a major gene for resistance to poplar leaf rust. Theoretical and Applied Genetics 103, 1129–1137

[11] Sun, H., D. Treco, N. P. Schultes, and J. W. Szostak. 1989. Double-strand breaks at an initiation site for meiotic gene conversion. Nature 338:87–90.

[12] Github repository for DMAP [http://github.com/davidmam/DMAP.git]

[13] A Golden Path file format specification [http://www.ncbi.nlm.nih.gov/projects/genome/assembly/agp/AGP_Specification.shtml]

[14] General Feature Format specification [http://www.sanger.ac.uk/resources/software/gff/spec.html]

[15] Stam P (1993) Construction of integrated genetic linkage maps by means of a new computer package: Join Map The Plant Journal 3, 739–744

